# Differential valuation and learning from social and non-social cues in Borderline Personality Disorder

**DOI:** 10.1101/305938

**Authors:** Sarah K Fineberg, Jacob Leavitt, Dylan S Stahl, Sharif Kronemer, Christopher D. Landry, Aaron Alexander-Bloch, Laurence T Hunt, Philip R Corlett

## Abstract

**Background:** Volatile interpersonal relationships are a core feature of Borderline Personality Disorder (BPD), and lead to devastating disruption of patients’ personal and professional lives. Quantitative models of social decision making and learning hold promise for defining the underlying mechanisms of this problem. In this study, we tested BPD and control subject weighting of social versus non-social information, and their learning about choices under stable and volatile conditions. We compared behavior using quantitative models.

**Methods:** Subjects (n=20 BPD, n=23 control) played an extended reward learning task with a partner (confederate) that requires learning about non-social and social cue reward probability (The Social Valuation Task). Task experience was measured using language metrics: explicit emotions/beliefs, talk about the confederate, and implicit distress (using the previously established marker self-referentiality). Subjects’ weighting of social and non-social cues was tested in mixed-effects regression models. Subjects’ learning rates under stable and volatile conditions were modelled (Rescorla-Wagner approach) and group x condition interactions tested.

**Results:** Compared to controls, BPD subject debriefings included more mentions of the confederate and less distress language. BPD subjects also weighted social cues more heavily, but had blunted learning responses to (non-social and social) volatility.

**Conclusions:** This is the first report of patient behavior in the Social Valuation Task. The results suggest that BPD subjects expect higher volatility than do controls. These findings lay the groundwork for a neuro-computational dissection of social and non-social belief updating in BPD, which holds promise for the development of novel clinical interventions that more directly target pathophysiology.

## Introduction

Learning whom to trust and when to revise trust attributions is a difficult but important task. People exhibiting extremes in trust can experience significant distress and personal risk, as in the very-low trust which characterizes paranoia (1) (3), and the very-high trust in Williams Syndrome (2) or amygdalar lesions (4). In Borderline Personality Disorder (BPD), trust is unstable, and interpersonal relationships involve recurrent episodes of rupture and repair. People with BPD suffer immensely, and attempt suicide at 50-fold the rate of the general population (5). Research investigating the mechanism of interpersonal problems in BPD is needed to identify targets for rational and effective treatment innovation. Low initial trust and rupture-promoting behavior in BPD have been modelled in the 10-round Trust Game, a brief economic exchange task with a partner (6). We aimed to extend those data by examining responses of people with BPD to instability of social and non-social information.

For this study, we used the Social Valuation Task (SVT), a laboratory-based reinforcement learning paradigm with social and non-social dimensions (7). The non-social dimensions are the color and the number on cards from which the subject chooses for potential points reward. The social dimension is advice from a confederate. Based on carefully computed contingencies, we can independently assess weighting of, and learning rates for, the social and non-social dimensions. Healthy undergraduates use both non-social and social dimensions (7), and they learn faster about each dimension when it is less reliable (8). FMRI dissociated social and non-social learning signals regionally (7).

Weighting of social versus non-social cues in the SVT in community samples correlates with self-reported traits. In healthy adults, self-reported autistic traits were directly correlated to poorer overall task performance, and inversely correlated to weighting social over non-social cues (9). Also, subjects with more autistic traits were worse at avoiding the influence of bad advice during the “volatile phase” of the task, when reward for the social cue was most unreliable. In another study, healthy women reported degree of psychopathic traits (10). As with autism score, the psychopathy sub-scale “social potency” – a measure of the ability to charm and manipulate others -- was inversely correlated with weighting of social cues, whereas “fearlessness” was inversely correlated with use of the non-social cue, and “stress immunity” was inversely correlated with weighting of both cue types. Also, Diaconescu et al. reported that in healthy men, self-reported stable social attributions correlated to stable beliefs about their game partner in a deception-free two-player version of the SVT (11). In sum, weighting of cues correlated with traits as expected. Autistic and manipulative traits correlated with decreased ability to make use of incoming social data, and stable social beliefs in self-reports correlated to stable social beliefs in the game.

We report here the first test of SVT behavior in a patient population. We model both weighting of social versus non-social cues and learning rates in response to changes in social and non-social reward volatility. Our main hypotheses about behavioral differences between BPD and control subjects were formulated in light of others’ work about BPD social experience, including interpersonal hypersensitivity, rejection sensitivity, and hyper-mentalization (12). We expected that people with BPD would be highly sensitive to small changes in the social environment (H1 - H3) and would change their behavior quickly in response to instability of the social environment (H4, H5):

H1. In BPD, social cues would be weighted more heavily than non-social cues.
H2. Social cues would be weighted more heavily by BPD than control subjects.
H3. Negative social cues would be weighted more heavily, and positive social cues would be weighted less heavily, by BPD than control subjects.
H4. Learning rate would increase more in during the volatile period in BPD than in control subjects.
H5. In BPD, learning rate would increase more in response to social volatility than non-social volatility.

We complemented our quantitative models of subject decisions with analysis of subjects’ experience. Consistent with our expectation that people with BPD would be more socially focused and responsive, especially to negative social data, we hypothesized that BPD subjects would express more surprise, suspicion, distress, and focus on the confederate. We expected that people (and BPD > control) would experience implicit distress due the periods of volatile and untrustworthy advice in the task, or after hearing at the end that the confederate was not in fact another player, and that we had intentionally misled them. We measured implicit distress by counting self-referential language (words like I, me, and mine), as they are known to increase with distress in mental and physical illnesses (13, 14) (15).

## Methods and Materials

### Ethics

This protocol was written and conducted in accordance with the Declaration of Helsinki and was approved by the Yale Institutional Review Board (HIC protocol 1211011104).

### Subjects

Women aged 18-65 were recruited from the community, and subjects identified who were either without psychiatric pathology or quite symptomatic with BPD (**Tables 1, 2**). See Supplementary Methods and Materials for details.

**Table 1.** Subject demographics. Note that some individuals in the BPD group were taking multiple psychiatric medications.

|  | Control | BPD |
| --- | --- | --- |
| **n** | 23 | 20 |
| **Age (yrs)** | 33.86 +/- 2.93 (18-60) | 35.80 +/- 2.91 (18-63) |
| **Education (yrs)** | 14.78 +/- 0.57 (10-19) | 14.20 +/- 0.54 (11-20) |
| **Ethnicity** |  |  |
| Asian | 13% | 10% |
| Black | 30% | 15% |
| Hispanic | 9% | 5% |
| White | 44% | 55% |
| Not Reported | 4% | 15% |
| **Taking psychiatric medications** | 0% | 45% |
| Anti-depressant | 0% | 15% |
| Mood Stabilizer | 0% | 25% |
| Anti-psychotic | 0% | 15% |
| Benzodiazepine | 0% | 10% |
| **Current Relationship** |  |  |
| None | 30% | 35% |
| In a relationship | 26% | 35% |
| Married | 13% | 10% |
| No answer | 31% | 20% |
| **Has children?** |  |  |
| Yes | 22% | 17% |
| No | 48% | 55% |
| No Answer | 30% | 28% |
| Benzodiazepine | 0% | 10% |
| **Current Work** |  |  |
| None | 26% | 45% |
| 0-20 hours/week | 13% | 15% |
| 20+ hours/week | 22% | 15% |
| In school | 30% | 10% |
| No Answer | 9% | 15% |

**Table 2.** Subject characteristics. Mean and standard error of the mean are displayed for the NAART (reading test), two different BPD self-reports (SCID2-BPD and BSL), depression self-report (BDI), and anxiety self-report (BAI). T-tests comparing control to BPD subject scores revealed no difference in reading score between groups, but significant differences in each of the self-reports.

|  | Control | BPD | p |
| --- | --- | --- | --- |
| **NAART** | 20.90 +/- 2.09 | 19.20 +/- 1.69 | 0.53 |
| **SCID2-BPD** | 0.90 +/- 0.28 | 9.00 +/- 0.76 | <0.001 |
| **BSL** | 5.95 +/- 1.60 | 33.00 +/- 4.29 | <0.001 |
| **BDI** | 3.10 +/- 1.06 | 21.40 +/- 2.95 | <0.001 |
| **BAI** | 7.75 +/- 2.32 | 23.00 +/- 2.86 | <0.001 |

### Self-report scales

Please refer to the Supplementary Methods and Materials for details.

### Social Valuation Task Design

The SVT was implemented as described by Behrens et al. (**Figure 1**) (7). For a detailed description of the task, please refer to the Supplementary Methods and Materials.

**Figure 1.**
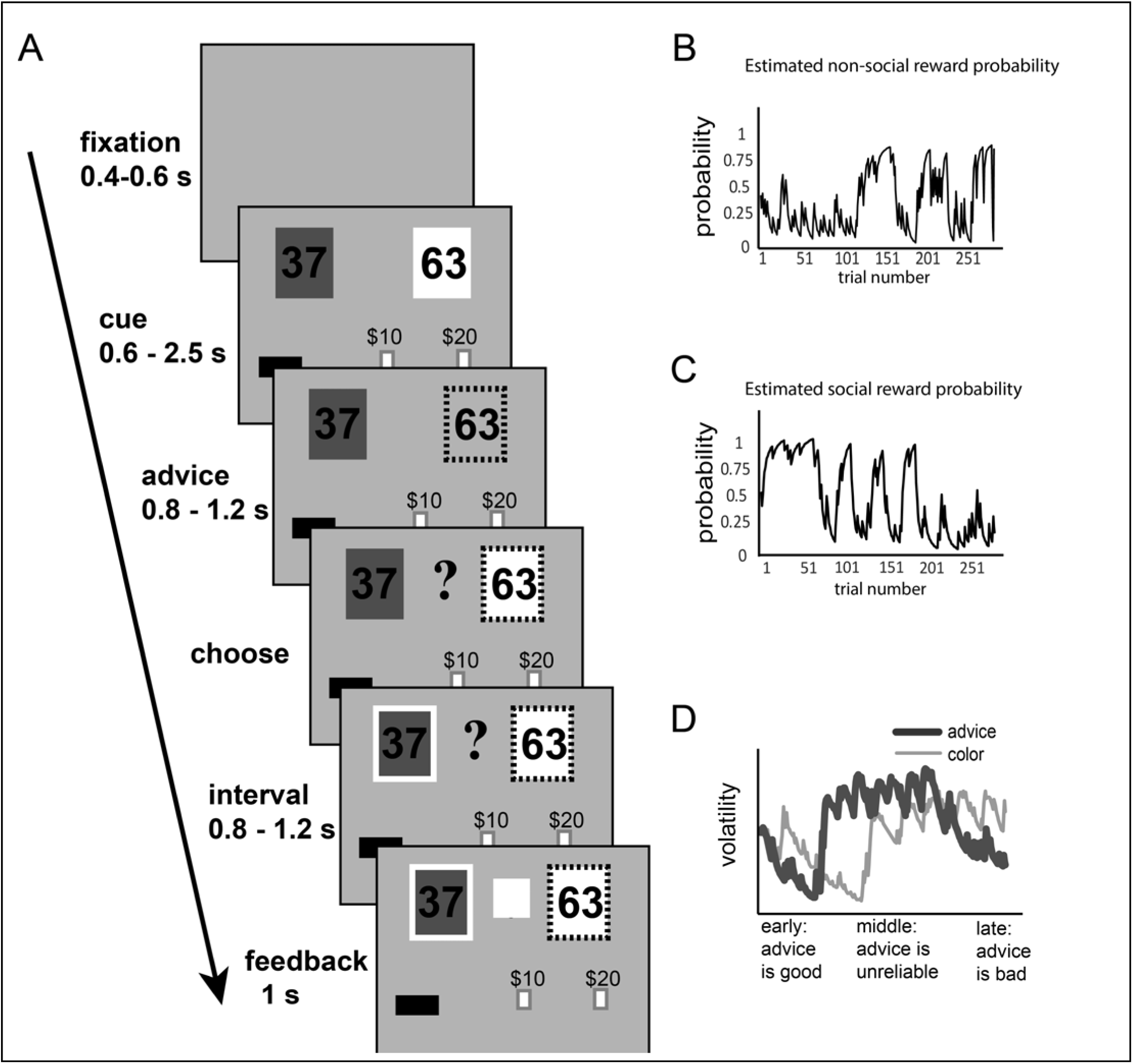
Task Design. A) Cartoon of task phases including elements displayed to the subject in each phase, and timings. Note that the subject makes her choice in the “choose” phase, and sees the correct answer in the “feedback phase”. These six phases are repeated in each of 290 trials. B) Changing probability of reward for the non-social cue (green) over time. C) Changing probability of reward for the social cue (advice) over time – note separate pattern from non-social cue. D) Changes in the volatility of the non-social (thin line) and social (thick line) reward probabilities represented as volatility over time. X-axis is again time (trial number), and quality of advice is indicated as it changes from low-volatility and trustworthy advice at the beginning through a period of high-volatility advice, to a period of low-volatility but untrustworthy advice at the end of the task.

### Confederate

The task confederates were 20 - 30 year old white women trained for consistent interaction with subjects and consistent performance during the demonstration task.

### Debriefing

Immediately after the task, subjects were audio-recorded talking with study staff in response to a list of specific questions and statements about the task experience. We asked 4 questions before the disclosure that the confederate was not actually a second game player, then 2 more questions after the disclosure. We examined the transcribed language from the debriefings. We counted the number of times that the subject mentioned the confederate.

To capture shifts in emotional state before versus after the disclosure, we examined the use of self-referential pronouns in subject speech (14). Transcribed speech was analyzed with Linguistic Inquiry and Word Count (LIWC) (16), which returns frequency of specific categories (we used “first-person pronouns”) as count/total words. We used repeated-measures ANOVA to test for interaction between time (before versus after disclosure) and group (BPD versus control).

### Modelling task behavior: relative cue weighting

Variables influencing subject decisions were examined with mixed models in the statistical program R using the package ‘lme4’ (17). Probability and volatility values were those derived by Behrens et al. from their Bayes-optimal model (8). Non-social variables were points (difference between point magnitude for green and point magnitude for blue), probability green is correct, and volatility of green being correct. Social variables were current trial advice, current advice weighted by probability advice is correct, current advice weighted by volatility of advice being correct, and refusing current advice after recent betrayal. We also tested a series of time windows on recent betrayal (incorrect advice) or help (correct advice): within x trials, where x = 1, 3, 4, 5, 6, or 7. Each variable was centered and z-scored to facilitate comparison of coefficients across factors. The impact of clinical group was tested separately for each of the above predictor variables, v (modelled as fixed effects in the mixed models). Likelihood ratio tests were used to compare nested models. For details of model comparisons, please refer to the Supplementary Methods and Materials section.

### Modelling task behavior: learning rates

Subject learning rates were modelled using the R package “hBayesDM” and the function “bandit2arm_delta” using default parameters (18). This package calculates mean learning rates for each subject based on the Rescorla-Wagner (delta) equation. See details in the Supplementary Methods and Materials.

The SVT has two phases for the non-social cue: stable (trials 1 – 130) and volatile (trials 131 – 290) (**Figure 1B, D**). There are three phases for the social cue: stable reliable (trials 1-70), volatile (trials 71 – 210), and stable unreliable (trials 211 – 290) (**Figure 1 C, D**). In our analysis, learning for each cue was modelled based on the trials in each phase. Repeated-measures ANOVA was used to test for group x phase interaction for each cue.

## Results

### Demographics

Subjects (control n = 23, BPD n = 20) were matched on age (t = −0.47, df = 39, p = 0.641) and education (for years in school t = 0.751, df = 39, p = 0.457, for reading score t = 0.631, df = 39, p = 0.53) (**Table 1**). BPD subjects had significantly more severe BPD, depression, and anxiety (**Table 2**). All the subjects were able to complete the task, and their final point scores did not differ by group or symptom burden (**Figure 2A**).

**Figure 2.**
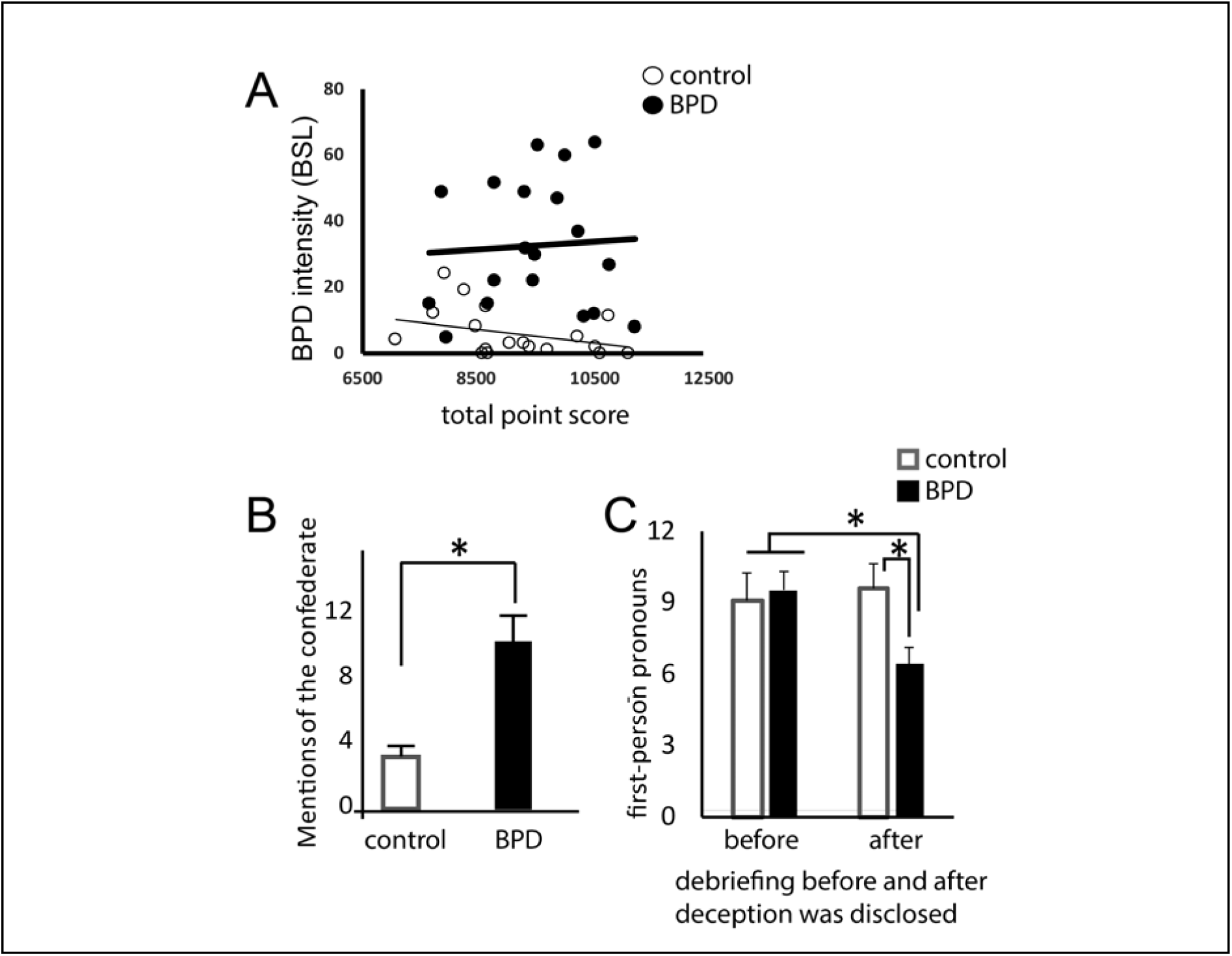
Task Experience. A) BPD and control subjects achieve similar final point scores in the game (t-test p = 0.345), and final point score does not correlate with BPD symptom score in either group (Pearson correlation Control r = −0.352, p = 0.140, BPD r = 0.061, p = 0.804). **B)** In the post-task debriefing, BPD subjects talk significantly more about the confederate than do control subjects. Error bars represent standard error of the mean (t-test p = 0.01). **C)** In the post-task debriefing, BPD and control subjects refer to themselves with similar frequency before the deception is revealed. However, after they hear that the computer, not the confederate, was providing the advice, self-referential language is significantly less in BPD than in control subjects. In a repeated-measures ANOVA, there was a trend-level difference by time (F = 3.03, p = 0.09) and a significant difference for the time x group interaction (F = 6.16, p = 0.02).

### BPD patients talk more about the confederate, but show lower implicit distress in response to task

As a preliminary test for enhanced focus on social cues in BPD, we counted references to the confederate in audio recordings of the post-task debriefing. There were more mentions of the confederate in BPD versus control subject language (**Figure 2B,** mean BPD = 11.60, SE = 2.34, mean control = 4.77, SE = 1.34; t = −2.67, df = 21, p = 0.01). The two groups did not differ in expressed surprise (mean BPD = 0.8, SE = 0.29; mean control = 0.77, SE = 0.23; t= −0.08, df = 21, p = 0.93), distress (mean BPD = 0, SE = 0; mean control = 0.15, SE = 0.10, t = 1.3, df = 21, p = 0.21), or suspicion (mean BPD = 0.10, SE = 0.10; mean control = 0.08, SE = 0.08, t = −0.19, df = 21, p = 0.85).

Though none of the subjects demonstrated overt distress during or after the task, we also tested for implicit distress. We used a previously established language measure: frequency of self-referential words (see Introduction). We analyzed subject language before and after we revealed the deception (that the social cues were controlled by the computer, not the human confederate). Control subjects used similar levels of self-referential words before and after disclosure. BPD subjects used similar levels to controls before disclosure, but significantly fewer afterward (**Figure 2C,** time x group interaction F = 6.16, p = 0.02). This suggests that in BPD subjects, distress actually <u>decreased</u> after the deception was revealed.

### H1/2: People with BPD weighted social more heavily than non-social cues

We tested the impact of non-social and social cues on subject choices in the SVT (**Figure 3**). To test our first and second hypotheses, we examined the weights of non-social and social cues in subject decisions. Each of the variables was a significant contributor to subject decisions, and contributed differently to decisions between groups. Specifically, BPD participants were more likely than controls to choose green when reward probability was higher (**3A**, likelihood ratio chi-square statistic = −4.03, p =0.045, reward probability coefficient = 0.41, group coefficient = 0.11) and less likely than controls to choose green when likelihood of reward became more volatile (**3C** likelihood ratio chi-square statistic = −3.48, trend-level significance p = 0.062, reward volatility coefficient = −0.17, group coefficient = 0.10). They were also more likely than controls to choose green if the difference between points for green and points for blue was larger (**3E,** likelihood ratio chi-square statistic = −4.07, p = 0.044, Δpoints coefficient = 0.30, group coefficient = 0.11). BPD participants were more likely to go with the advice if reward probability was higher compared to controls (**3B,** likelihood ratio chi-square statistic = −5.98, p = 0.015, social reward probability coefficient = 0.46, group coefficient = 0.12).

**Figure 3.**
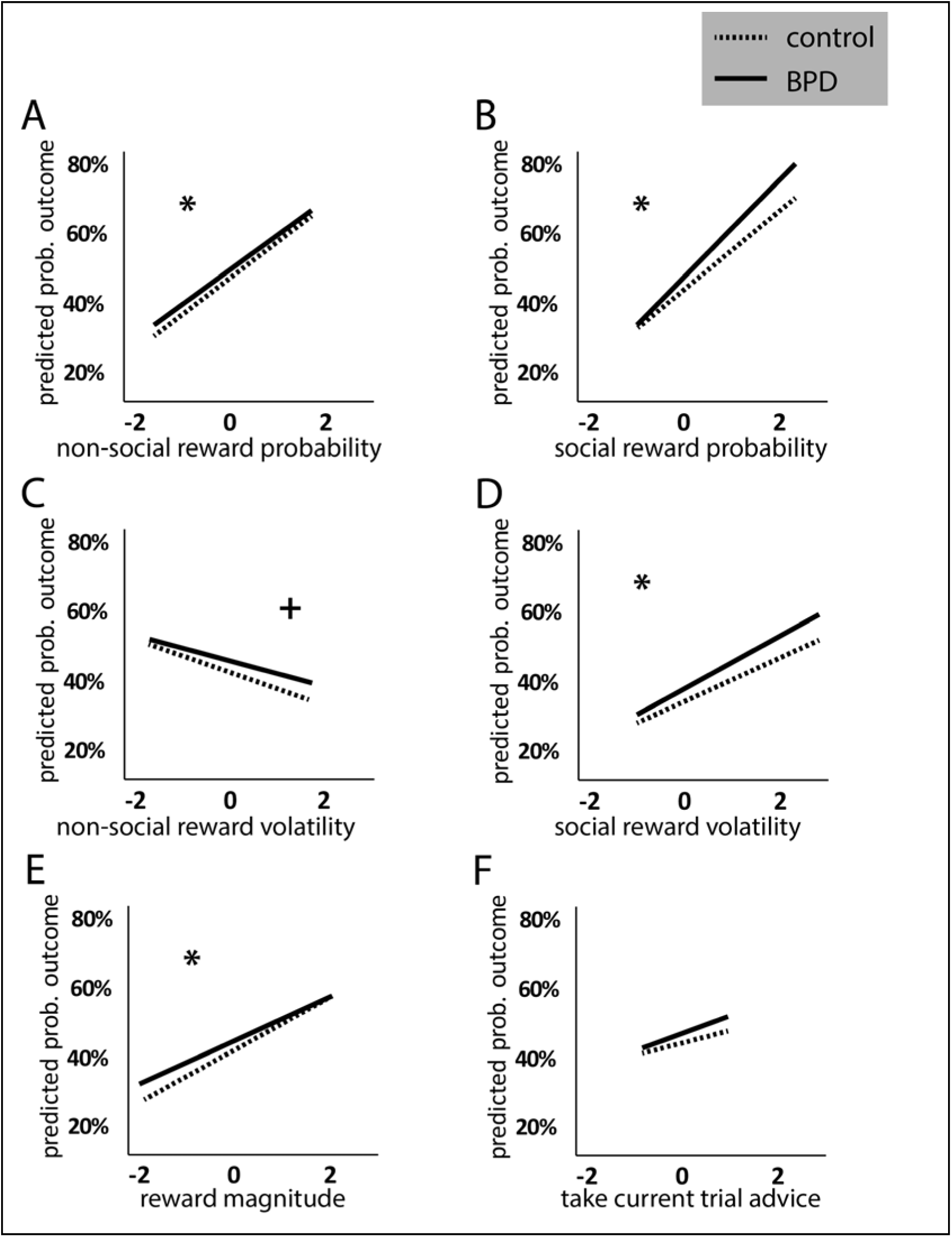
BPD and control subjects use cues differently to make decisions. Mixed-effect logistic regression models were run to measure the effect of non-social and social predictors on subject choices. Predicted probability of choosing green is plotted over z-scored values of each predictor. Non-social predictors (A,C,E) and social predictors (B, D) both performed differently between groups (except F, current trial advice). Significant Chi-square for models with versus without group term at p < 0.05 is indicated by *. Trending Chi-square for models with versus without group term at p < 0.1 is indicated by +.

Of interest, and perhaps surprising, is that both groups (BPD > control) were also <u>more</u> likely to take the advice when social reward likelihood was more volatile (**3D**, likelihood ratio chi-square statistic = −4.96, p = 0.026, social volatility coefficient = 0.21, group coefficient = 0.12). However, the model describing outcomes predicted by group and current trial advice (what Behrens et al. termed “blindly following advice”) did not detect statistical differences (**3F**). In sum, we found that people with BPD made significant use of both non-social and social cues. Interestingly, people with BPD weighted both social <u>**and non-social** cues more heavily than controls, although</u> between-group differences were larger for weighting of social reward probability than for non-social reward probability (based on magnitude of difference between regression lines; as noted chi-square tests were significant in both cases).

### H3: People with BPD used positive social cues more than negative social cues to make decisions

To better understand the responses of BPD and control subjects to social cues in this interactive context, we next tested the predicted choices after recent betrayal (bad advice) or help (good advice) (**Figure 4**). We found a significant decrease in BPD patients in use of betrayal to avoid bad choices (**4A,** BPD> control, for betrayal within the last three trials, likelihood ratio chi-square statistic = −4.25, p= 0.039) and a trend toward a group x predictor interaction for increased use of help to find good choices in BPD patients (**4B, for help within the last 3 trials,** BPD > control, likelihood ratio chi-square statistic for group x predictor interaction = −3.79, p = 0.051).

**Figure 4:**
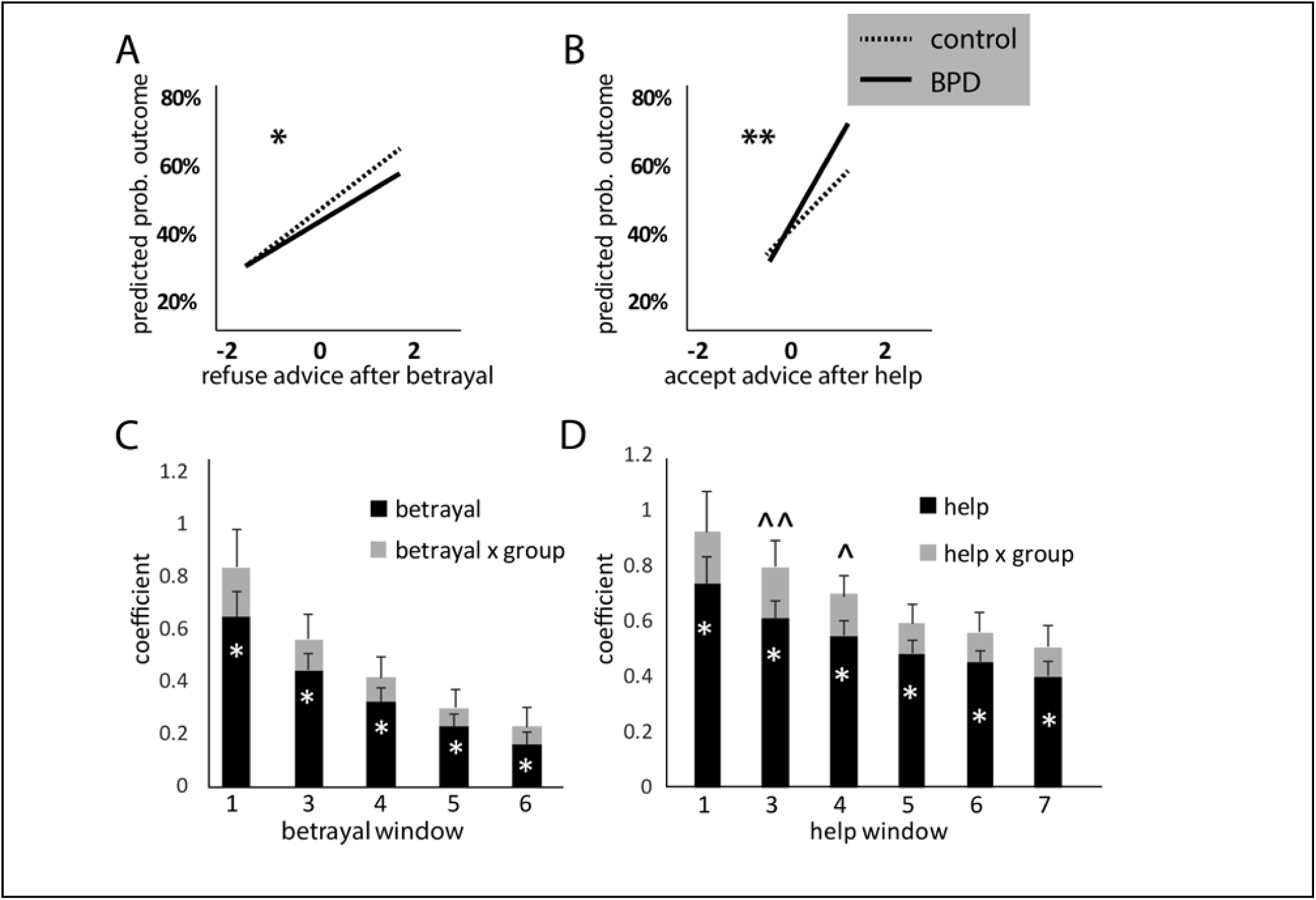
Both betrayal and help impact differently on BPD and control subjects. Predicted probability of outcome differed in BPD versus control subjects exposed to a recent instance of betrayal (incorrect advice) or help (correct advice). BPD subjects were more likely to refuse the advice after betrayal (**A,** * indicates significant effect of group at p < 0.05) and take the advice after help (**B,** ** indicates significant group * predictor effect at p < 0.05). **C** and **D** depict the results of a closer look at the use of betrayal and help over time. The bar graphs show regression results for a series of time windows, each with at least one betrayal (**C**) or help (**D**) event in the marked number of trials. For example, help window 1 means the advice was correct on the previous trial, help window 4 means the advice was correct on at least one of the previous 4 trials. * predictor p < 0.05, ^^ predictor x group p < 0.05, ^ predictor x group p < 0.1. All subjects showed diminishing use of these social cues with enlarging time windows; the significant group x predictor interaction in help windows 3 and 4 but not in betrayal windows suggests a slower decrement of help use for decision making in the BPD group compared to the control group. This is not an effect that we observed for the use of betrayal.

We also tested the rate of decrement in weighting of recent help or betrayal. As expected, both groups used help or betrayal less as the window size expanded (**Figure 4C, 4D**), but both help and betrayal were significant predictors of outcome out to at least a 7-trial window. However, it was help (the <u>positive</u> social cue) not betrayal (the <u>negative</u> social cue) that showed a trend towards a group x predictor interaction (use of help decayed more slowly in the BPD than the control group).

### H4/5: Learning rates reveal blunted response to increased reward volatility in people with BPD

We modelled learning rates for non-social and social rewards during the stable and volatile phases of the task. We found that control and BPD subjects learned at similar low rates about non-social data during the initial phase when reward probability was stable (**Figure 5A,** t = −1.38, p > 0.05). However, when reward probability became volatile (phase 2), control subjects increased their learning rate more than twice as much as did the BPD subjects (**Figure 5A,** significant group x condition interaction: F = 19.78, p < 0.001). Learning from social cues was slower in BPD than in controls during all three phases of the task (**Figure 5B,** stable reliable t = 4.02, p < 0.001, volatile t = 2.90, p < 0.01, stable misleading t = 3.44, p < 0.005), and response to volatility in BPD was significantly lower than in controls (**Figure 5B,** group x condition interaction, F = 5.81, p < 0.01). These results were surprising: we had hypothesized faster learning rates in BPD in response to reward volatility, and instead, we observed blunted response compared to controls for both non-social and social cues.

**Figure 5.**
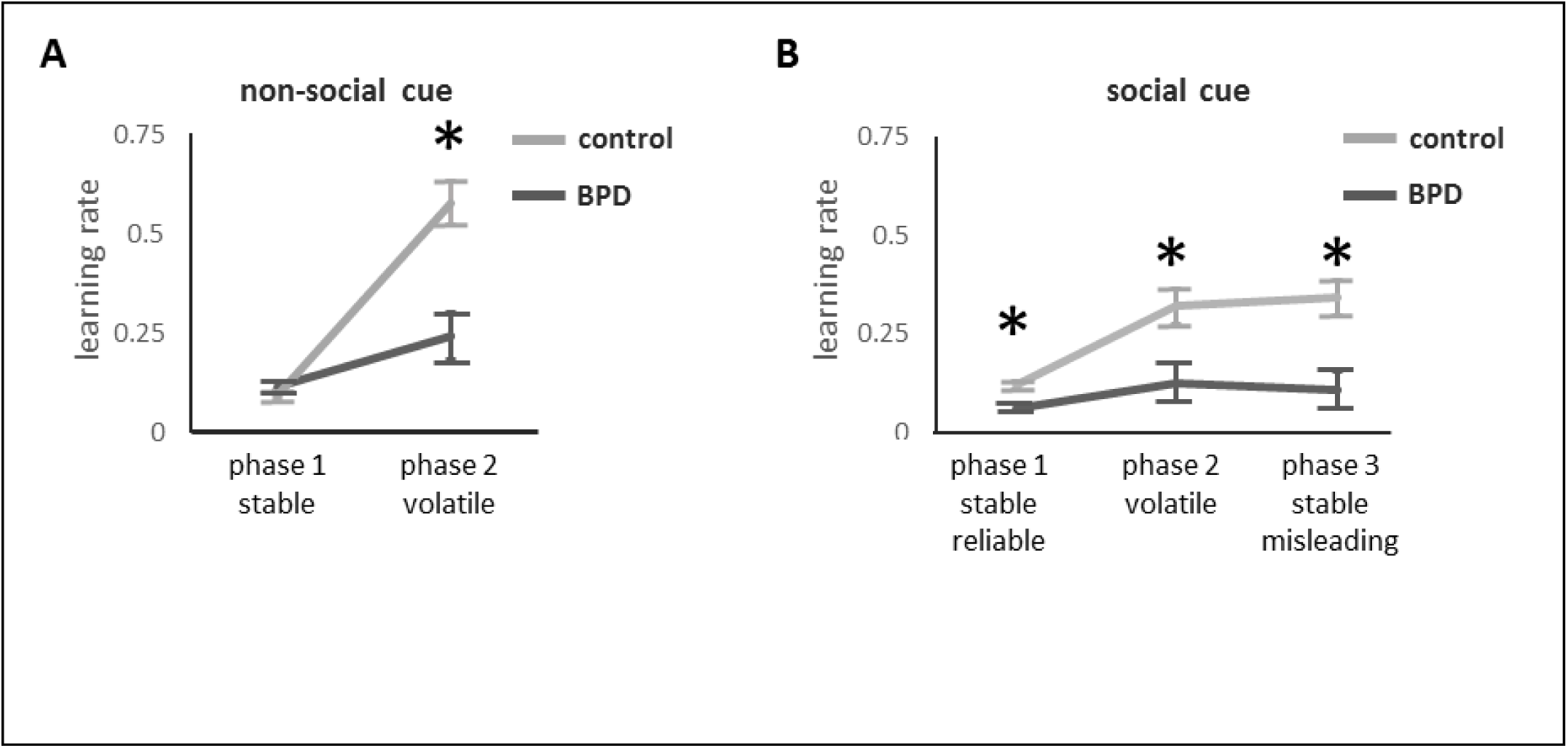
Learning rates were estimated for each individual differences analysed by group and condition. * indicates significant between-group difference. **A)** For learning from the non-social cue (color), there was a group x condition interaction (F = 19.78, p < 0.001, error df = 1, hypothesis df = 41). Learning was slow and not significantly different by group when reward probability was stable (t = −1.38, p > 0.05). However, when reward probability became volatile, the BPD group had a blunted response (less of an increase in learning) compared to controls (t = 4.12, p < 0.001). **B)** For learning from the social cue, a group x condition interaction was also observed (F = 5.81, p < 0.01, hypothesis df = 2, error df = 40). Here, learning was slower in the BPD group than in the control under all three conditions: stable reliable t = 4.02, p < 0.001, volatile t = 2.90, p < 0.01, stable misleading t = 3.44, p < 0.005), and there was again a blunted response to volatility of reward probability under the volatile condition.

## Discussion

In this first study of the SVT in a patient population, we examined task performance in people with BPD, a condition defined by prominent interpersonal problems. In this extended interactive paradigm, women with BPD did indeed focus on social experience, weighting social over non-social cues to make decisions. However, we also found that a negative social experience (incorrect advice) was a less potent and less durable influence on subject choice than a positive social experience (correct advice).

Previous work in non-interactive paradigms, such as Reading the Mind in the Eyes and morphed face challenges has identified a strong negative attribution bias (reviewed in (19, 20)). BPD patients attend quickly to negative faces and spend more time looking at them. A small number of studies have tested the response of BPD patients to social interaction games using brief paradigms. In the 10-round “Trust Game”, players with BPD responded with low initial trust and failure to coax defecting partners back to play (6). A key difference between the “Trust Game” paradigm and the SVT is that the latter combines social and non-social cues; negative social experiences in the SVT may therefore lead subjects to use non-social information more prominently in guiding their choices, which is not possible in the “Trust Game”.

Unlike the reported problems in task performance with increased sub-clinical autism and psychopathy symptoms, our sample of women with BPD completed the SVT with final point scores similar to controls, and used both social and non-social cues to make decisions. Non-social <u>and</u> social cues were weighted more heavily in the BPD group than in the control group. This may suggest that people with BPD are more attentive to <u>all</u> of the cues around them. Previous work describing learning in BPD has had mixed results. Work in brief non-social paradigms found that BPD state (emotional arousal) but not traits (BSL score) predicted problems with learning acquisition and vice versa for reversal learning (21). Others found no difference in reversal learning (22), but deficits in working memory in BPD (23).

In the extended and more complex social interaction in the SVT, we were able to examine not only low-probability but also low-reliability social rewards. In contrast to the defection (failure to coax) that was observed in the trust game after low-payoff trials (6), we saw increased use of social cue under conditions of high social reward volatility here (in both groups, but BPD > control), as if subjects were redoubling their efforts to remain socially engaged.

We extended previous reports of the effect of personality/mental health traits on SVT behavior by examining learning rates. We replicated the Behrens et al. report that control subject learning rates increase under conditions of reward volatility. However, we were surprised to observe that BPD subjects showed only half the learning rate increase that control subjects did in response to non-social reward volatility, and barely responded at all to social reward volatility. One possibility is that the BPD subjects assume higher baseline volatility of all environments and contingencies, such that high volatility is not surprising, and does not prompt updating. This is consistent with research demonstrating early life adversity, especially neglect (= volatility) is a key risk factor for BPD. Our observation that people with BPD decrease their self-referential language after confirmation of deception is consistent with this idea. The BPD subjects used fewer language markers that connote distress once they were informed of the deception, consistent with the idea that they harbor assumptions that the social world is unreliable. For someone sensitive to volatility, attending closely to cues, but not updating rapidly may be adaptive. Future work could model prior beliefs about social and non-social cues to test our hypothesis that people with BPD assume higher baseline volatility.

This insensitive learning account of BPD social behavior is also consistent with a recent report describing a computational model of BPD trustee responses in the 10-round Trust Game (24). They had previously found that people with BPD fail to coax a defecting partner to re-engage in economic exchange. They now report that a hierarchical belief model significantly benefits from parameters describing a player’s own irritability and beliefs about the partner’s irritability. Here, irritability means the likelihood of retaliation on a low economic offer. BPD players were significantly less sensitive to their partners’ irritability than control players, and the authors suggest that this leads to missed opportunities to respond: a player doesn’t coax if she doesn’t detect early cues that the partner is becoming irritable or is likely to disengage.

Particular strengths of this work include the use of a patient population, in fact, we carefully screened participants for non-clinical status (controls) or significant symptoms (DIB score > 8 in the BPD group). Our subjects met and interacted with the confederate before starting the task, perhaps increasing their ability to engage with the task in a manner that reflects their real-world social behavior. The Social Valuation Task is a lengthy interactive task that combines non-social (one’s own beliefs) with social (others’ counsel) for decision-making at each trial. The task architecture includes orthogonal periods of volatility for the non-social and social cues which allows us to model the relative use of each data source in decision-making.

There are several limitations of this work. The sample is small and all female. We would expect gender differences in the expression of BPD, and cannot generalize these results to men. We mostly see our very symptomatic patient group as a strength of this study, but our exclusion of people with fewer or less intense symptoms does preclude dimensional analysis of the impact of symptom burden on behavior. Also, we did not test what aspect of psychopathology (e.g. negative affect, anxiety, negative attribution bias) is most correlated to the outcome measures. Future work will benefit from a larger sample with psychopathology control groups (depression, anxiety, PTSD), or a dimensional approach with subjects who vary in intensity of core symptoms, relevant co-morbidities, and treatment history. Symptoms also fluctuate widely with time in BPD: future work should test the relationship of task behavior to subject emotional state by self-reports (e.g. Positive and Negative Affect Scale) or physiologic arousal (e.g. galvanic skin response or pupillary dilation). Conversely, testing the impact of the task on subject state could further validate our finding of distress in subjects’ post-task language. To examine changes in task behavior over time, as we might wish to do to understand how subject learning changes with treatment in clinical practice, we will need interactive tasks that do not rely on deception and are more likely to work in repeated measures, as in an SVT-inspired task developed by Diaconescu et al. (11).

Future work should also examine social learning in BPD in settings that engage more realistic, and real-world, relationships. Schilbach et al. have scored tasks based on whether the subject is interactive (versus passive) and engaged (versus dispassionate), with the truly interactive engaged task representing a more informative approach to social cognitive experiments, “second person neuroscience” (25). In this frame, emotional engagement is tightly linked to bodily experience and the complex and immediate dynamics of perceiving another’s action states. In our experiment, subjects do meet and briefly interact with their game partner (confederate) before the task. The SVT asks the subject to make decisions using the partner’s advice. The subject also knows that she and the game partner are incentivized to adjust their choices to the way the other behaves. Therefore, we see this task as interactive – the subject is playing the game in cooperation (competition?) with her game partner (or at least she thinks so). However, in terms of emotional engagement, the subject does not see or hear the game partner during the task, so no emotion is conveyed through bodily movements or language. Though some might argue that people with BPD are often surprisingly engaged based on little data, we think that this task likely does not meet the Schilbach criteria for an engaged task, and that future work could test this axis by including more possibility for engagement.

Our approach to modelling also has some limitations. We used mixed-effects regression models to compare our subjects’ behavior to an ideal Bayesian observer (a model without random effects). There is a potential tension here in terms of how much we account for individual variation in the processes that generate subjects’ behavior. As discussed above, we aim in the future to model additional parameters, such as subjects’ prior beliefs. In this initial effort to extend the work of Behrens et al., we adhered closely to their approach. However, Diaconescu et al. have used the Hierarchical Gaussian Filter (HGF), allowing individual differences and obtaining subject-specific estimates of approximate Bayesian inference (9, 11, 26). Modelling patient behavior using the HGF may also detect subtler between-group differences (11).

Computational psychiatry, which focuses on the development of mechanistic models linking clinical symptoms to neurobiology and observed behavior through computational parameters, has already begun to describe the neurobiology of learning under volatile conditions (26), and to help mental health researchers improve prognostic prediction (27) and to make plans to more precisely target therapeutics (28). A computational psychiatry of social interaction holds promise for honing existing treatments and building new ones in BPD and other disorders with prominent interpersonal symptomatology (29).

## Acknowledgements

We would like to thank our research participants. Several members of our lab helped as confederates and with recruitment efforts: Emily Finn, Carol Gianessi, Kristen Budde, Erin Feeney, Margot Reed, Sasha Deutsch-Link, Megan Ichinose, Taylor McGuiness, and Rachel Zubi. Clinicians at the Yale New Haven Hospital Intensive Outpatient Program and in the Connecticut Mental Health Center Ambulatory Services also helped with subject recruitment. Megan Ichinose, Lindsey Conkey, Mary Zanarini, and Emily Ansell gave very helpful advice on research methods.

This work was supported by the Connecticut State Department of Mental Health and Addiction Services through the Connecticut Mental Health Center Clinical Neuroscience Research Unit. Sarah K. Fineberg was supported by NIMH Grant no. 5T32MH019961, “Clinical Neuroscience Research Training in Psychiatry” and a NARSAD Young Investigator Award (2014-2016). Dylan S. Stahl was supported by the Richter Memorial Fund and the NSF COAST Award as an undergraduate student at Knox College. Aaron Alexander-Bloch was supported by was supported by NIMH Grant no. 5T32MH019961, “Clinical Neuroscience Research Training in Psychiatry”. Laurence Hunt was supported by a Sir Henry Wellcome Fellowship from the Wellcome Trust (098830/Z/12/Z), a NARSAD Young Investigator Award, and by the NIHR Oxford Health Biomedical Research Centre. Philip R. Corlett was funded by an IMHRO/Janssen Rising Star Translational Research Award and CTSA Grant Number UL1TR000142 from the National Center for Research Resources (NCRR) and the National Center for Advancing Translational Science (NCATS), components of the National Institutes of Health (NIH), and NIH roadmap for Medical Research. The contents are solely the responsibility of the authors and do not necessarily represent the official view of NIH, the NHS, NIHR or the UK Department of Health.

Other results from other experiments conducted in this patient sample have been published elsewhere (https://www.ncbi.nlm.nih.gov/pubmed/29248760). Preliminary versions of these results have been shown as a poster and an oral presentation at the National Association for the Study of Personality Disorders and as a poster at the Society of Biological Psychiatry. A pre-print is available on bioarxiv: https://www.biorxiv.org/content/early/2018/04/22/305938.

The authors report no conflicts of interest or financial disclosures.

